# Deep Brain Stimulation induced normalization of the human functional connectome in Parkinson’s Disease

**DOI:** 10.1101/537712

**Authors:** Andreas Horn, Gregor Wenzel, Friederike Irmen, Julius Hübl, Ningfei Li, Wolf-Julian Neumann, Patricia Krause, Georg Bohner, Michael Scheel, Andrea A. Kühn

**Affiliations:** Department of Neurology, Movement Disorders and Neuromodulation Section, Charité – University Medicine Berlin; Berlin School of Mind and Brain, Humboldt-Universität zu Berlin; Department of Neuroradiology, Charité – University Medicine Berlin

**Author notes:** Corresponding Author **Material & Correspondence:** Andreas Horn, MD, PhD, Charité – University Medicine Berlin, Charitéplatz 1, 10117 Berlin.

## Abstract

Neuroimaging has seen a paradigm shift from a formal description of local activity patterns toward studying distributed brain networks. The recently defined framework of the ‘human connectome’ allows to globally analyse parts of the brain and their interconnections. Deep brain stimulation (DBS) is an invasive therapy for patients with severe movement disorders aiming to retune abnormal brain network activity by local high frequency stimulation of the basal ganglia. Beyond clinical utility, DBS represents a powerful research platform to study functional connectomics and the modulation of distributed brain networks in the human brain. We acquired resting-state functional MRI in twenty Parkinson’s disease (PD) patients with subthalamic DBS switched ON and OFF. An age-matched control cohort of fifteen subjects was acquired from an open data repository. DBS lead placement in the subthalamic nucleus (STN) was localized using a state-of-the art pipeline that involved brain shift correction, multispectral image registration and use of a precise subcortical atlas. Based on a realistic 3D model of the electrode and surrounding anatomy, the amount of *local* impact of DBS was estimated using a finite element method approach. On a *global* level, average connectivity increases and decreases throughout the brain were estimated by contrasting ON and OFF DBS scans on a voxel-wise graph comprising eight thousand nodes. Local impact of DBS on the motor STN explained half the variance in global connectivity increases within the motor network (R = 0.711, p < 0.001). Moreover, local impact of DBS on the motor STN could explain the degree of how much voxel-wise average brain connectivity normalized toward healthy controls (R = 0.713, p < 0.001). Finally, a network based statistics analysis revealed that DBS attenuated specific couplings that are known to be pathological in PD. Namely, coupling between motor thalamus and motor cortex was increased and striatal coupling with cerebellum, external pallidum and STN was decreased by DBS.

Our results show that rs-fMRI may be acquired in DBS ON and OFF conditions on clinical MRI hardware and that data is useful to gain additional insight into how DBS modulates the functional connectome of the human brain. We demonstrate that effective DBS increases overall connectivity in the motor network, normalizes the network profile toward healthy controls and specifically strengthens thalamo-cortical connectivity while reducing striatal control over basal ganglia and cerebellar structures.

## Introduction

Deep Brain Stimulation (DBS) is a highly efficacious and established treatment option for Parkinson’s Disease (PD; Deuschl *et al.*, 2006) and a multitude of mechanisms of action have been proposed over the years (Lozano and Lipsman, 2013). In recent years, these underwent a paradigm shift away from localized stimulation of specific areas toward global modulation of distributed brain networks (Akram *et al.*, 2017; Horn *et al.*, 2017; Litvak *et al.*, 2011; McIntyre and Hahn, 2010; Vanegas Arroyave *et al.*, 2016). To elucidate such network effects of DBS, modern-day neuroimaging methods become increasingly meaningful (Horn, 2019). In addition, DBS could reversely be a powerful tool to study network changes as a function of precisely targeted stimuli.

One of the few neuroimaging options to study the functional organization of the brain is functional magnetic resonance imaging (fMRI). Intrinsic associations between subparts of the brain may be estimated using resting-state (rs-) fMRI and in this way, the “functional connectome” of the brain may be explored (Biswal *et al.*, 2010). When blood oxygenated level dependent (BOLD) signals of two brain regions are correlated over time, these have been called functionally “connected” in the literature (Fox *et al.*, 2005; Friston, 2011), although this measure includes highly indirect connections (Varoquaux and Craddock, 2013). Until recently, it was not straight-forward to acquire rs-fMRI data in patients with DBS implants, let alone with the stimulator switched on in the scanner. The reason was that no official certificate of device manufacturers allowed this practice and only limited pioneering work by a few specialized centers – the Jech and Foltynie groups should be mentioned among others – investigated changes of fMRI data under DBS in a so far limited fashion (Jech *et al.*, 2001; Joshua Kahan *et al.*, 2014); supplementary table S1. In a first study involving 4 patients, Jech *et al*. showed that the BOLD signals increase in ipsilateral subcortical structures under DBS (Jech *et al.*, 2001). In a case-report, Stefurak *et al*. then showed more distributed signal increases in (pre-)motor cortices, ventrolateral thalamus, putamen and cerebellum under effective DBS (Stefurak *et al.*, 2003). Seminal work by Kahan *et al*. in 2014 described changes of direct, indirect and hyperdirect pathways of the basal ganglia – cortical loops under DBS using dynamic causal modeling (Joshua Kahan *et al.*, 2014). The same data was used to fit a computational mean-field model that was able to identify additional potential DBS targets beyond the classical STN target used to treat Parkinson‘s Disease (PD; Saenger *et al.*, 2017). In a formal literature analysis, we identified further studies that used fMRI under active DBS in humans and animal models so far (table S1). In summary, STN-DBS in PD may lead to increased overall functional connectivity in the premotor cortex (Mueller *et al.*, 2013) and strengthened cortico-striatal and thalamo-cortical pathways in fMRI (Joshua Kahan *et al.*, 2014).

**Table 1:**
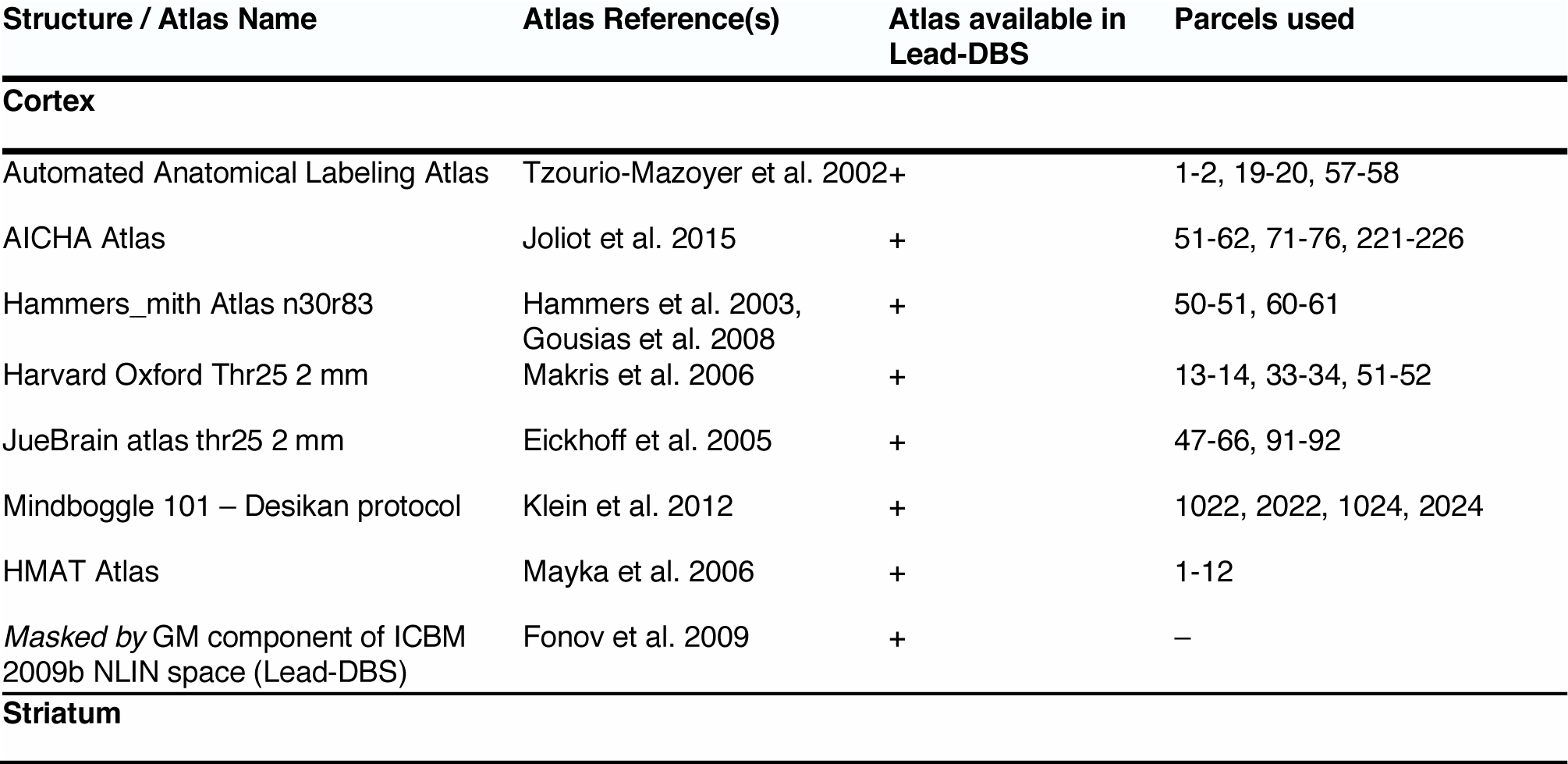
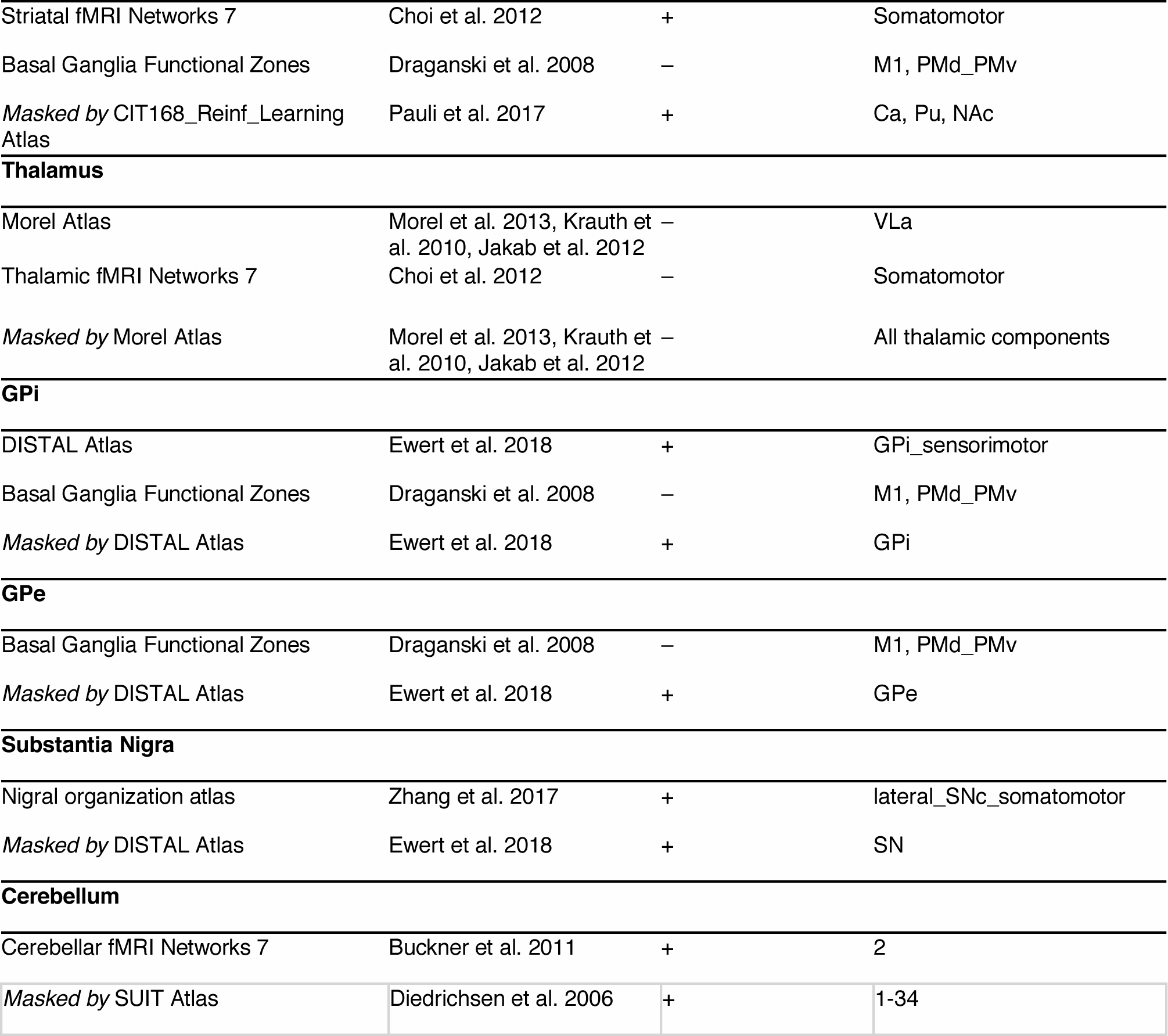
Atlases that contributed to motor network definition.

In sum, even though there was no official allowance for active DBS in the fMRI until recently (see methods), the concept was explored in small case numbers. One issue that has been neglected in prior studies is that slight changes of millimeters in lead placement lead to large differences in clinical improvement (Horn *et al.*, 2019) and similarly, slight differences in connectivity profiles of DBS electrodes may be used to predict clinical improvement across patients, cohorts and DBS centers (Horn *et al.*, 2017). Finally, small variations in lead placement even explain changes of behavior in motor (Neumann *et al.*, 2018) and cognitive (Irmen *et al.*, 2018) domains. Thus, we argue that it is crucial to incorporate DBS lead placement into an analysis of their impact on distributed brain networks. Instead, prior studies characterized fMRI changes in DBS ON vs. OFF contrasts and, by doing so, implicitly assumed that the DBS effect was equal in each patient. In reality, the impact of each DBS electrode on the target structure varies across patients and may be used as a regressor to better explain network changes.

In the present study, we investigated a cohort of 20 PD patients at rest under DBS ON and OFF conditions. We characterized changes in average connectivity (i.e. centrality) of brain regions and laid special focus on network changes *as a function of* the degree of motor STN-DBS modulation. Based on minor differences in DBS electrode placement, different amounts of motor STN volume were stimulated in each patient. As a result, we expected correspondingly differing changes in motor cortical activation that should be stronger or weaker as a function of electrode placement. Moreover, because differences in therapeutic effects are equally dependent on lead placement (Horn *et al.*, 2017; 2019), we expected network properties to normalize toward healthy controls depending on lead placement. We expected that an optimally placed lead would result in strong modulations in the motor network, normalizing toward the network properties found in healthy controls. In contrast, poorly placed leads would not result in strong motor network changes.

## Methods

### Literature analysis

To summarize previous studies in the still novel field of DBS-fMRI, a systematic medline literature analysis was conducted using the search-string (“Deep brain stimulation” OR “DBS”) AND (“Parkinson’s disease” OR “Parkinson” OR “Parkinsonian” OR “thalam*” OR “pallid*” OR “subthalamic”) AND (“functional connectivity” OR “resting-state fMRI” OR “fMRI”). This resulted in n = 64 studies on August 31, 2018. Predefined inclusion criteria for studies were: (i) investigation of effects of DBS in subcortical targets such as thalamus, pallidum or subthalamic nucleus in PD patients or (potentially healthy) animals using fMRI; (ii) bilateral/unilateral stimulation of DBS target during fMRI scan. Exclusion criteria were: (i) studies reporting safety of fMRI in DBS patients or animals without reporting structures affected by DBS; (ii) resting-state fMRI studies in healthy humans; (iii) studies assessing medication effects without DBS; (iv) meta-analyses and reviews. Relevant references were identified by title and abstract (n = 17) and further judged based on inclusion and exclusion criteria, originality, quality and actuality of the study resulting in n = 15 remaining papers. Another 7 publications were added by screening reference lists of included articles. Results are summarized in table S1.

### Patients and imaging

#### DBS cohort

Twenty patients suffering from Parkinson’s Disease that underwent DBS with target STN were included in the present study. Inclusion criteria were at least > 3 months of active STN-DBS, an MR-conditional DBS system (in agreement with the Medtronic whole-body CE certificate; Medtronic, 2015), no metal implants beside the DBS system, no largely predominant affective or cognitive non-motor symptoms of the disease, no excessive tremor at rest that would make scanning impossible. The study was carried out in accordance with the Declaration of Helsinki and was approved by the internal review board of Charité – Universitätsmedizin Berlin (master vote # EA2/138/15). All patients had undergone DBS surgery for idiopathic PD between April 2010 and April 2016 and received 2 quadripolar DBS electrodes (model 3389; Medtronic, Minneapolis, MN). Before surgery, patients received preoperative MRI and neuropsychological testing to exclude structural or psychiatric comorbidities. During surgery, microelectrode recordings were performed to verify lead placement. Clinical variables, including age, sex, L-dopa response and L-dopa equivalent dose (LEDD) at baseline were recorded. Postoperative improvement scores were recorded 12 months after surgery under Med OFF and DBS ON/OFF conditions using clinical DBS stimulation parameters.

After multiple studies had shown fMRI acquisition with implanted DBS devices to be safe (Carmichael *et al.*, 2007; Jech *et al.*, 2001; Sutton *et al.*, 2008), and elaborate modeling and animal testing by the company Medtronic, their Activa® portfolio received an extended MR conditional CE certificate for full-body MRI in 2015 (FDA approval followed 2016). Under this extended certificate, it became officially feasible to acquire fMRI data under active DBS in humans (Medtronic, 2015). Limits, such as a maximal scanning time of 30 minutes and a B1+RMS value below 2.0 μT apply – and it is only allowed to stimulate using bipolar settings. Thus, postoperatively, patients were scanned in a Siemens Magnetom Aera 1.5T MRI. After a structural scan, a resting-state fMRI scan of ∼9 minutes was carried out in DBS ON condition. Patients were then briefly taken out of the scanner to turn the impulse generator off. An experienced neurologist (either GW or JH) waited for symptoms to reappear. After that, the same scan was repeated (DBS OFF condition). Scan parameters were as follows: T1 MP-RAGE: Voxel-size 1 × 1 × 1 mm, TR 2200 ms, TE 2.63 ms. Rs-fMRI EPI scan: Voxel-size 3 × 3 × 3 mm, 24 slices with distance factor 30%, A >> P phase encoding, TR 2690 ms, TE 40 ms, FoV readout 200 mm. 210 volumes were acquired for each condition (total scan length of rs-fMRI scans 2 × 9.42 min). A brief dMRI scan was acquired before the rs-fMRI scans but was not used in the present study. Total scan time was close to but below 30 minutes and the B1+RMS value was kept below 2.0 uT at all times (conforming to the MR-conditional regulations of the Medtronic Activa CE-certificate).

#### Normative control subjects

Structural and rs-fMRI data from 15 control subjects were obtained from the Parkinson’s Disease Progression Marker Initiative (PPMI; ppmi-info.org; Marek *et al.*, 2011) database. These were processed in the same way as patients of the present study using Lead connectome software (Horn and Blankenburg, 2016; Horn *et al.*, 2014) and had been made available in form of a normative group connectome in (Horn *et al.*, 2017). Thus, details in acquisition and processing scheme are described elsewhere (Horn *et al.*, 2017; Marek *et al.*, 2011).

### Electrode localization and modeling the volume of tissue activated

All data was processed in Lead-DBS software (www.lead-dbs.org; Horn and Kühn, 2015) using the enhanced default workflow of version 2.1.7 described in (Horn *et al.*, 2019). Within the pipeline, Lead-DBS uses a multitude of open source tools (see below and Horn *et al.*, 2019 for details). Briefly, all (pre- and postoperative) MRIs were linearly co-registered to the preoperative T1 anchor modality using SPM (https://www.fil.ion.ucl.ac.uk/spm/software/spm12/; Friston *et al.*, 2004). Registration between post- and preoperative T1 was further refined using the “brainshift correction” module in Lead-DBS which focuses on the subcortical target ROI (Schönecker *et al.*, 2009) and thus minimizes nonlinear bias introduced by opening the skull during surgery. Data was normalized into standard space (ICBM 2009b NLIN ASYM; (Fonov *et al.*, 2009); henceforth referred to as “MNI space”) in a multispectral fashion (i.e. using T1- and T2-weighted) in parallel. This was done with the symmetric diffeomorphic registration algorithm implemented in Advanced Normalization Tools (http://stnava.github.io/ANTs/; Avants *et al.*, 2008) using the “effective (low variance)” preset with subcortical refinement as implemented in Lead-DBS (Horn *et al.*, 2019). This exact normalization method was recently evaluated best-performing in a large comparative study (Ewert, Horn, *et al.*, 2018) and STN definitions obtained by the method rivaled those of manual expert segmentations in precision. Electrode trajectories and contacts were automatically pre-localized and manually refined using Lead-DBS. Based on the (bipolar) stimulation settings active in the scanner, the electric fields (E-Fields) around the electrode were modeled using a finite element approach defined in (Horn *et al.*, 2017; 2019). Briefly, an adapted version of the FieldTrip/SimBio pipeline that is part of Lead-DBS (http://www.fieldtriptoolbox.org/; https://www.mrt.uni-jena.de/simbio/; Vorwerk *et al.*, 2018) was used which uses a four-compartment anatomical model of the electrodes and surrounding tissue. All analyses were performed using the impact of the E-Field on the STN but were repeated with a binarized version of it, the VTA that is a more commonly applied model in the field. To approximate the VTA, the E-field gradient magnitude was thresholded at a heuristic value of 0.2 V/mm. This value has been frequently been used in similar context (Åström *et al.*, 2009; 2014; Horn *et al.*, 2017; Vasques *et al.*, 2009).

### Estimating the impact of DBS on motor STN

As mentioned above, in the DBS context, the VTA has most often been modeled in binary form (Åström *et al.*, 2014; Butson *et al.*, 2007; Dembek *et al.*, 2017; Horn *et al.*, 2017; McIntyre and Grill, 2002). However, a recent study showed superior results when directly using the E-field magnitude (instead of a binarized version; Horn *et al.*, 2019). Building upon this, here, we estimated the factor *impact of DBS on the motor STN* by multiplying the (unthresholded) E-field gradient magnitude map with the atlas-defined mask of the motor STN (Ewert, Plettig, *et al.*, 2018) and summed up voxels of the resulting image. Intuitively, this grasps the amount of voltage gradient that is present within the motor STN. Of note, all results reported in the present study would hold true with similar (and significant) effect sizes when repeating analyses using a binarized VTA, instead.

### Definition of a (sensori-)motor network parcellation

To estimate changes in the motor network of key regions within the basal ganglia cerebellar cortical loop, sensorimotor functional zones of cortex, striatum, thalamus, GPi/GPe, substantia nigra and cerebellum were defined based on multiple brain atlases available to the authors. Table 1 summarizes which atlas components were used for this parcellation. If multiple atlases defined a parcel, definitions were summed up to include voxels defined by every atlas available. Finally, for each component, parcels were masked by a definition of the whole structure (e.g. the motor thalamus parcel was masked by a thalamic mask) which is also denoted in table 1. Code that generates the final atlas is available online (www.github.com/leaddbs/leaddbs). Throughout the manuscript, we refer to this network as “motor” network, but it does include premotor and sensory domains.

### Estimating changes in rs-fMRI based connectivity

rs-fMRI scans under DBS ON and OFF conditions were preprocessed using Lead-Connectome (www.lead-connectome.org; Horn *et al.*, 2014). The pipeline is described in detail elsewhere (Horn, 2015; Horn and Blankenburg, 2016; Horn *et al.*, 2014) and largely follows the suggestions outlined in (Weissenbacher *et al.*, 2009). Briefly, rs-fMRI time series were detrended and motion parameters, as well as mean signals from white-matter and cerebrospinal fluid were added as nuisance regressors for noise removal. No global signal regression was performed. Signals were then band pass filtered between 0.009 and 0.08 Hz. Finally, spatial smoothing with an isotropic 6 mm full-width half maximum Gaussian kernel was applied. As all tools involved in the present study, the preprocessing pipeline is openly available to ensure reproducibility (https://github.com/leaddbs/leaddbs/blob/master/connectomics/ea_preprocess_fmri.m).

A voxel-wise parcellation scheme with 8k nodes in grey matter (see “Voxelwise parcellations (Lead-DBS)” under http://www.lead-dbs.org/helpsupport/knowledge-base/atlasesresources/cortical-atlas-parcellations-mni-space) was inversely registered to rs-fMRI volumes (via the T1 anchor modality) and signals were sampled from each node. Marginal correlations between time series were estimated, leading to two 8k × 8k adjacency matrices for DBS ON and OFF conditions in each subject. Based on these, average connectivity between each node and all other 8k – 1 nodes were calculated. Formally, this measure may be referred to as strength centrality (the weighted version of degree centrality; (Rubinov and Sporns, 2010)). However, we will henceforth refer to it as *average connectivity* since this seems a fitting intuitive term for what we calculate. Likewise, *changes in average connectivity* denote the difference between DBS ON and OFF conditions. Spearman correlations between local DBS impact and fMRI-derived measures were calculated. Random permutation (× 5000) was conducted to obtain P-values.

Subsequently, changes in connectivity between specific parts of the “motor network” (table 1) as a function of motor STN-DBS were estimated using the network based statistics (NBS) approach as implemented in the GraphVar toolbox (http://www.rfmri.org/GraphVar; Kruschwitz *et al.*, 2015). This involved nonparametric testing of DBS ON/OFF differences in each link that were explained by the degree of motor STN-DBS stimulation. Overall significance was then estimated using the NBS approach on the whole graph matrix (Zalesky *et al.*, 2010).

## Results

Patients (N = 20) were 63.7 ±6.6 (STD) years old at time of surgery and the sample included 4 women. Healthy control subjects were 59.5 ±11.9 years old and 4 of 15 subjects were female.

Patients were scanned 30 ±21 months after surgery with a minimum of 4 months in one subject. All other subjects had been implanted at least 11 months at the time of the scan. Due to logistic reasons (see methods), patients were scanned in Med ON condition and first scanned in DBS ON followed by the DBS OFF condition (with a brief interval of 5-15 minutes in which the impulse generator was turned off).

Before surgery, patients had a Levodopa response of 36.8 ±11 % (percent improvement in Med ON vs. OFF conditions; from 35.9 ±7.6 to 18.9 ±6.8 points on the UPDRS-III scale; lacking scores of two patients). Levodopa-equivalent dose of medication was 724.7 ±441 mg (lacking data of one patient). Under effective DBS, motor symptoms improved by 43 ±18 % (from 38.8 ±10.9 points before in DBS OFF to 22.0 ±9.8 in DBS ON; lacking scores of two patients). Please note that stimulation parameters during rs-fMRI differed since they had to be in bipolar setting and were not all taken at the same time of the 12 months postoperative evaluation. The aim was to best match unipolar long-term stimulation settings with the bipolar settings used during rs-fMRI acquisition. Still, for logistic reasons, a second UPDRS score could not be taken at the time of scanning and thus a matching UPDRS-III improvement for the DBS applied in the scanner is not available in this cohort.

Electrodes were all placed in the subthalamic region (fig. 1) although five patients had an overlap between their bilateral volume of tissue activated (VTA) and the motor parts of the bilateral STN (as defined by the DISTAL atlas; Ewert, Plettig, *et al.*, 2018) below 4 mm^3^. In comparison, the average coverage of motor STN stimulation in the rest of the sample was 36.4 ±20.5 mm^3^ which amounts to 50.6 % of the bilateral nucleus volume. Congruently, clinical improvements in this subgroup of patients were significantly worse compared to the rest of the cohort (29.5 ±14 % vs. 49.4 ±17 % improvement on the UPDRS-III scale; p = 0.039) – although as mentioned above VTAs used for clinical assessment and during the rs-fMRI experiment were not directly comparable.

**Figure 1:**
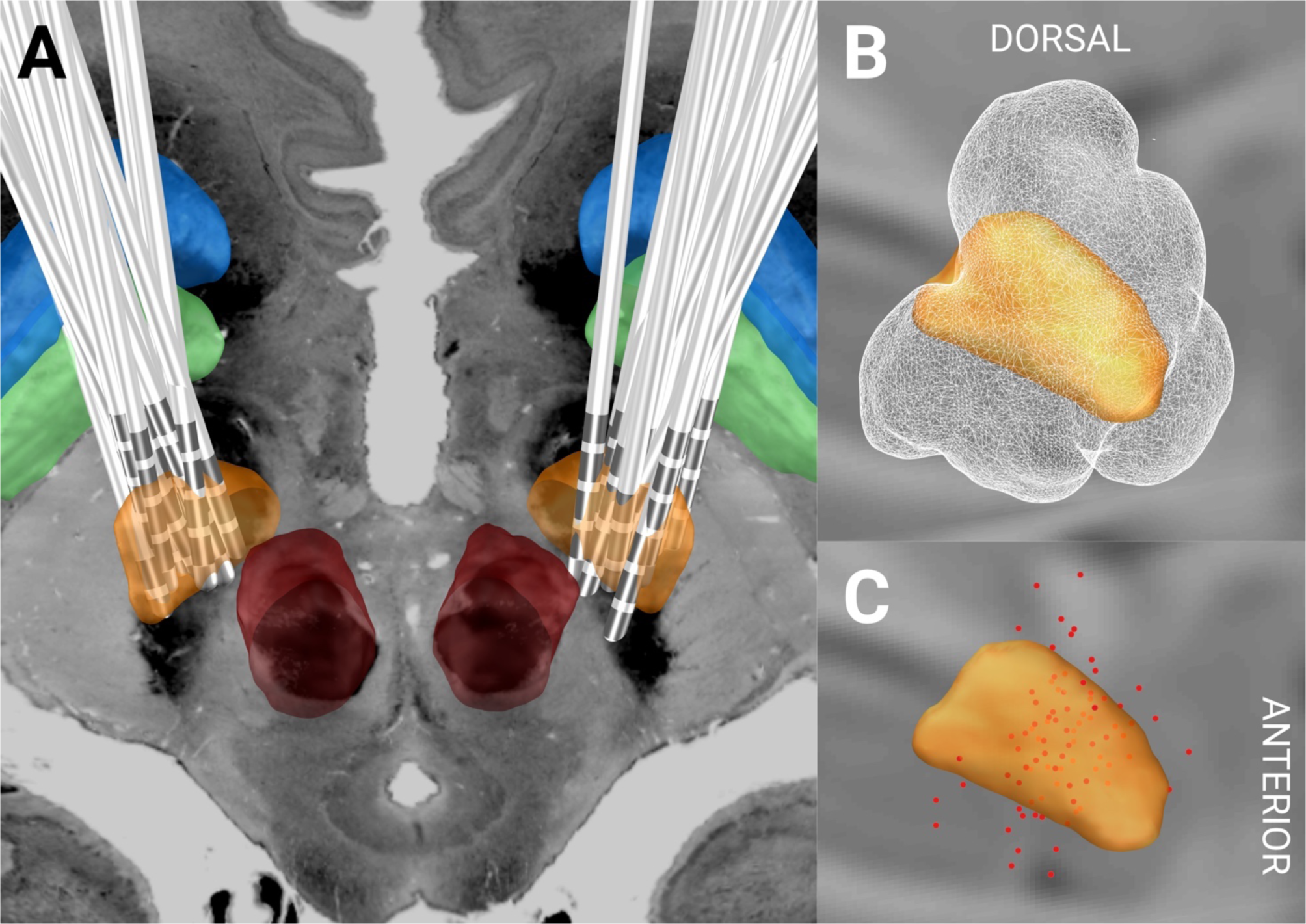
DBS electrode placement in the sample of 20 patients. A) 3D lead reconstructions together with STN (orange), GPi (green), GPe (blue) and red nucleus (red) as defined by the DISTAL atlas (Ewert, Plettig, *et al.*, 2018). An axial section of the BigBrain dataset (Amunts *et al.*, 2013) at z = −10 mm is shown. Right hemispheric volumes of activated tissue (B) and active (bipolar) stimulation contacts (C) were nonlinearly flipped to the left hemisphere to show the combined stimulation volume of the group (white wireframes) and the active contacts (red dots) in relationship to the STN.

To investigate whether subject motion in the scanner differed between DBS ON and OFF conditions, motion parameters were estimated using SPM12 and framewise displacements calculated following the approach of Power *et al*. (Power *et al.*, 2014; fig. 2 C). Average displacement values for DBS ON and OFF conditions were not significantly different (0.26 ±0.17 mm in ON vs. 0.30 ±0.17 mm in OFF, p = 0.32) and within a typical range of tolerable movement (Ardekani *et al.*, 2001).

**Figure 2:**
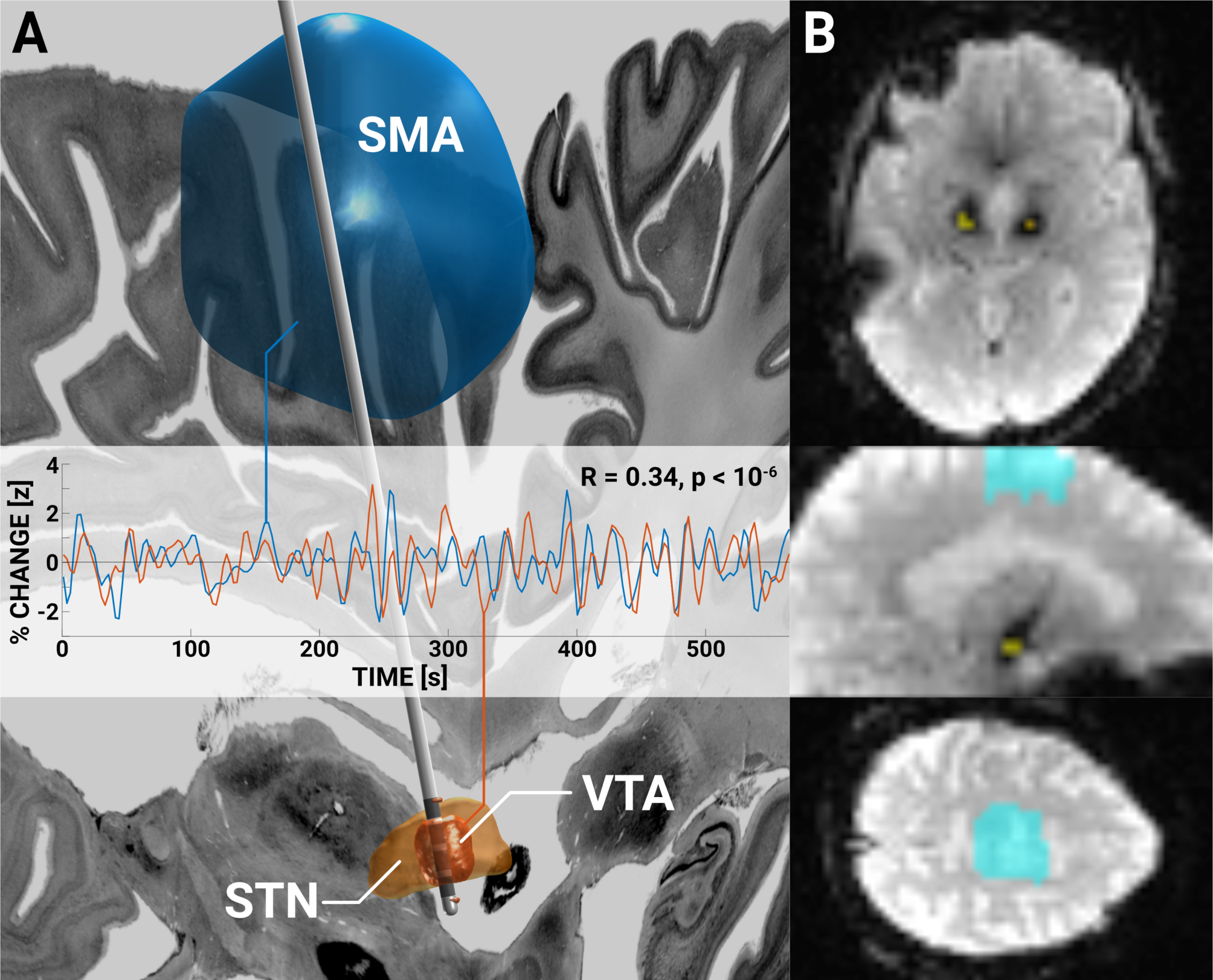
A-B) Resting state time series sampled from the left VTA (red/yellow) and bilateral SMA (blue/cyan) in an example patient in DBS ON condition. Despite presence of electrode artifacts, useful BOLD time series could be sampled from the vicinity of the electrode (i.e. the VTA) that show functional connectivity to cortical motor regions (such as the SMA). A) shows a 3D reconstruction and example time series from the left hemisphere SMA. B) shows the actual data with ROI overlays. Susceptibility artifact of the lead extension can be seen on the axial slice. In this patient, correlation between the two contralateral VTA time series was R = 0.41, correlation between left/right VTA and SMA were R = 0.34 / 0.25. In the DBS OFF condition, values were similar (left to right VTA: R = 0.34; left/right VTA to SMA: R = 0.32 / 0.42). Sagittal and coronal sections of the BigBrain dataset (Amunts *et al.*, 2013) shown as backdrops.

To investigate whether BOLD time series sampled directly from the region of the electrode / volume of tissue activated (VTA) could be used despite the prevalence of metal artifacts, we first calculated functional connectivity between bilateral VTAs and second between VTAs and supplementary motor area (SMA) in DBS OFF conditions. The SMA was chosen given its functional role and positive connectivity to the STN (Akram *et al.*, 2018; Horn *et al.*, 2017). In some patients, correlation values of R > 0.3 were prevalent between bilateral VTAs, suggesting the STN time series around the electrode are useable but variability in lead placement introduced strong variance across patients (average absolute R values were 0.16 ±0.11). Crucially, the connectivity between the two electrodes could be explained by lead placement: The more similar their impact on the motor STN, the higher their connectivity (R = 0.45 at p < 0.05).

Functional connectivity values between bilateral VTAs and SMA were similar, R-values ranged up to 0.24 (average absolute R values were 0.14 ±0.06). Here, lead placement did not significantly correlate with connectivity between VTA and SMA (R = 0.31, p = 0.09).

Figure 2 illustrates BOLD time series sampled from the VTA and the supplementary motor region (SMA) of a representative patient and gives a visual impression about the BOLD signal sampled directly at and around the electrode artifact.

Despite the apparent usefulness of the BOLD signal directly at the stimulation sites, the first and main analysis of this manuscript did *not* use this signal but rather investigated whole-brain connectivity changes within the motor network (cortex, cerebellum and basal ganglia). Average connectivity changes between DBS ON and OFF conditions were estimated on a graph of 8k nodes that covered the whole brain but were then sampled within the motor network (see table 1 for a proper definition of the network). In other words, overall connectivity changes between each pair of voxels in the whole brain was defined and changes in the motor network were then correlated with impact of DBS on the motor STN. Across subjects, changes in average connectivity of the motor network were correlated with the impact of the DBS electrode on the motor STN (R = 0.71, p < 0.001). Figure 3 shows these results across the group and highlights two example subjects. In the first one (pt. #9, fig. 3 top), overlap between the VTA and motor STN is large – i.e. it represents a case with (near) optimally placed DBS electrodes. As expected, in this patient under DBS, average connectivity increases strongly – and predominantly in the motor network. The contrary is the case in a patient where the leads are largely *outside* the STN where as expected little to no modulation of the motor network takes place (pt. #1, fig. 3 bottom). Across the group, this relationship between DBS placement and the connectivity increase in the motor network was strong. In fact, 50% of the observed variance in motor network changes (which are exclusively informed by rs-fMRI) may be explained purely by lead placement (which is exclusively informed by structural MRI).

**Figure 3:**
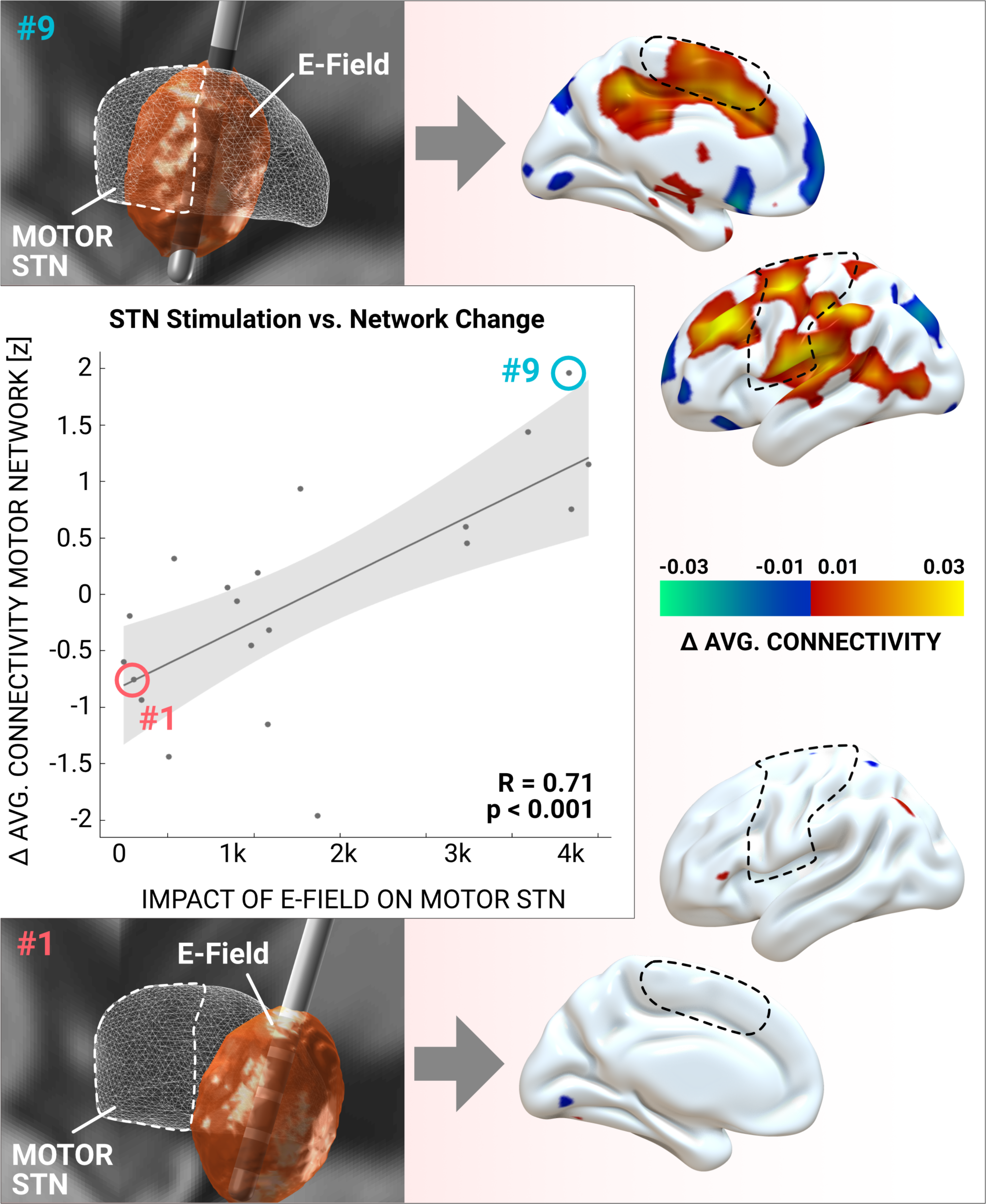
The amount of stimulated motor STN explains connectivity changes within the motor network across patients. The larger the weighted overlap between E-Field and motor STN, the more connected the motor network becomes (on average to all other brain regions). Here, “motor network” refers to sensorimotor zones (including premotor domains) in cortex, cerebellum, striatum, pallidum and thalamus (visible parts outlined by dashed lines; see methods). Example subjects #1 and #9 are marked in the scatter plot and their electrode placement (left) and cortical changes in connectivity induced by DBS (right) are shown. Crucially, cortical connectivity changes were calculated independently from knowledge about electrode placement. Both E-Field and cortical changes are visualized for the left hemisphere only but were similar on the right hemisphere.

This analysis concluded that changes of average connectivity throughout the motor network were dependent how strongly the DBS electrode modulated the motor STN. This latter factor is dependent on electrode placement and the applied stimulation amplitudes (as known parameters) and on the electrode-tissue impedance (which was not measured and thus is an unknown parameter). To exclude the possibility that the impedance had a significant impact on the observed relationship, we performed a stochastic sensitivity analysis to account for the potential bias introduced by tissue impedance. Specifically, E-Fields were re-calculated, in each patient, after scaling the assumed tissue conductivity values by factors of 0.5, 0.75, 1, 1.5 and 2. Based on these additional E-Fields that were calculated using adapted conductivity values, a 5000-fold permutation test was run and correlations were recalculated for each permuted combination. This analysis showed high robustness of the relationship toward the potential influence of the unknown factor tissue-impedance. In fact, all combinations of VTAs under different impedance assumptions revealed constantly significant relationships, with the lowest R-value at 0.47 (p = 0.028), the highest at 0.75 (p < 0.001) and the average at R = 0.58 ±0.04 (p = 0.004 ±0.004).

After this analysis, we investigated relationships between motor STN stimulation and connectivity changes of specific regions within the motor network. In post-hoc uncorrected head-to-head comparisons, this relationship was significant for the cortical (R = 0.50, p = 0.024) and cerebellar (R = 0.51, p = 0.022) regions of interest (ROI) but not for remaining ROI (striatum, thalamus, GPi, GPe, substantia nigra) with a positive but non-significant R-value for all regions.

Subsequently, connectivity profiles seeding from bilateral VTAs in DBS ON and OFF conditions were compared to the same maps informed by a normative connectome of age-matched healthy controls (Horn *et al.*, 2017; Marek *et al.*, 2011). DBS ON and OFF maps (top right and left maps in figure 4) were each compared to healthy controls (top mid map) by means of voxel-wise spatial correlation (as a similarity metric). Here, seed connectivity maps in DBS ON condition were significantly more similar to the ones obtained when using rs-fMRI data of healthy controls both on a group level (bottom right and left scatter plots in figure 4) and single subject level (bottom mid bar plots). This analysis shows that the overall connectivity of the DBS electrode “normalizes” toward healthy controls under DBS. In a subsequent analysis, we compared whole-brain average connectivity estimates (voxel-wise strength centrality) changes based on DBS. These estimates became more similar to the ones obtained in healthy controls as a function of motor-STN stimulation. The same two patients (#9 and #1 shown in top and bottom rows of figure 5) are shown as examples. Here, the average connectivity metrics across the whole brain – and not just the motor network – became more similar to healthy controls as a clear function of motor STN stimulation. Similarly to the analysis in fig. 3, ∼50% of variance in how much the overall average connectivity “normalizes” toward healthy controls could be explained just based on DBS electrode placement. In other words, the more optimal a DBS electrode was placed (as measured by overlap with the motor STN), the more normal the overall functional connectivity became (as measured by similarity to healthy controls).

**Figure 4:**
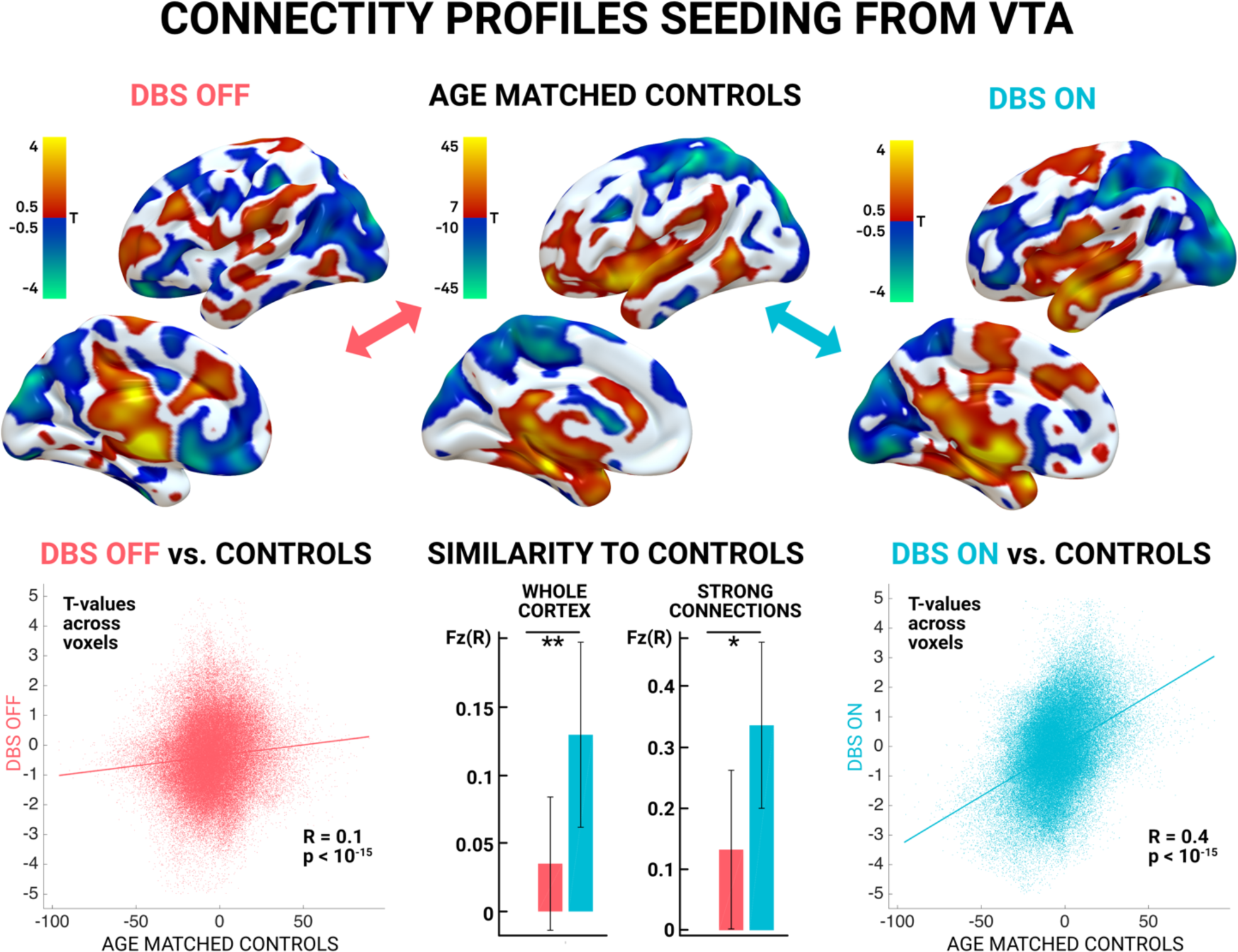
rs-fMRI functional connectivity profiles seeding from volumes of tissue activated (weighted sum of E-fields estimating the bilateral DBS stimulation volumes) to the rest of the brain. Patient specific data of DBS OFF (top left), ON (top right) conditions as well as maps derived from healthy age matched controls (bottom left) were compared. Unthresholded T-maps calculated across the group of 20 patients are shown in the top row. Spatial correlation values (that assess similarity) across cortical voxels between each DBS condition and the healthy controls were calculated on a group (scatter plots) and individual level (mid bottom) and were significantly higher between ON and healthy controls vs. OFF and healthy controls.

**Figure 5:**
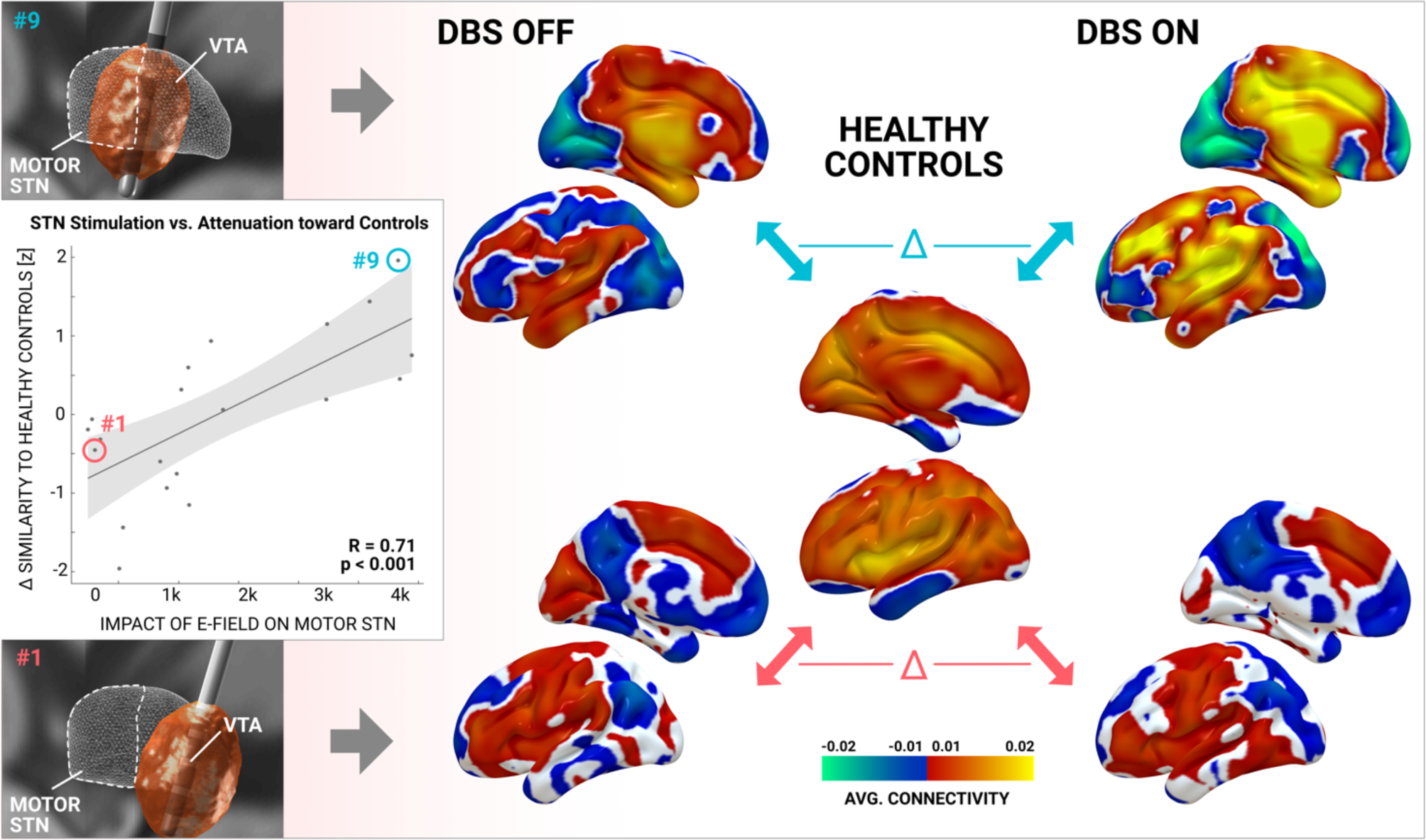
The shift of whole brain average connectivity profiles toward that of healthy controls (central) is dependent on the amount of motor STN stimulated (scatter plot left). Whole brain average connectivity profiles in DBS OFF (left) and ON (right) conditions of the same example patients as in fig. 3 are shown. Their connectivity distribution is spatially correlated with the one in healthy controls (to assess similarity) and the difference between conditions is related to the amount of motor STN stimulation in each patient. As can be seen in patient #9 (top row), DBS has a strong effect on the average connectivity profile. In contrast, profiles of patient #1 remain similar.

To further investigate exactly which connections would increase or decrease, we applied the network based statistics approach on DBS ON vs. OFF graph between ROI defined in table 1 (which include motor domains of STN, GPi, GPe, striatum, thalamus, cerebellum, substantia nigra and cortex). Again, changes were investigated as a function of motor STN DBS (weighted overlap between E-Field and motor STN). As expected, this showed a significant increase between thalamus and cortex and decreases between Striatum and Cerebellum, Striatum and STN as well as STN and GPe (fig. 6).

**Figure 6:**
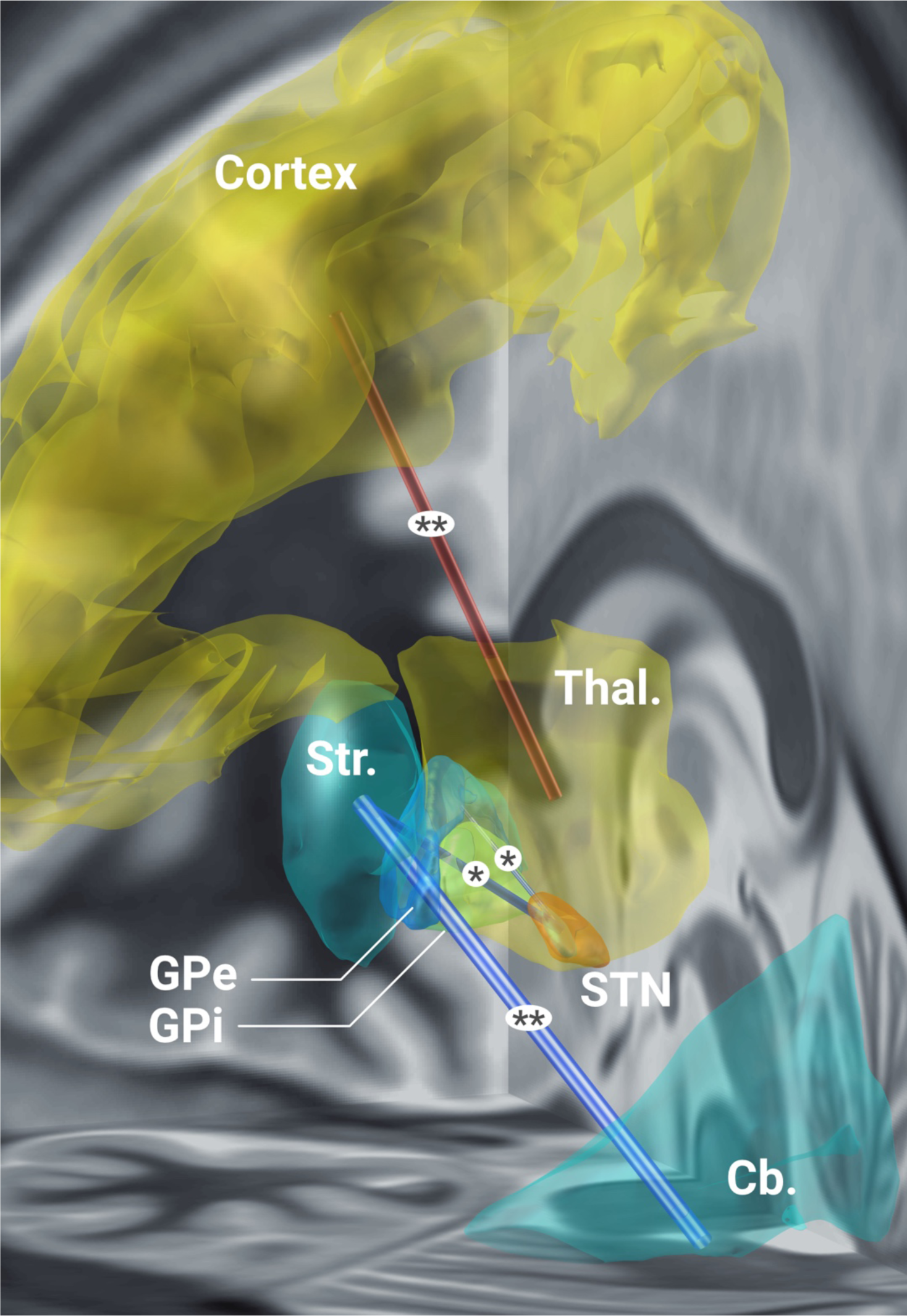
Specific connections in the motor network modulated by effective STN-DBS. Functional connectivity between motor thalamus and cortex (SMA, M1, S1) increase as a function of DBS impact on the motor STN. Instead, connectivity between motor striatum and cerebellum, motor striatum and motor STN as well as motor STN and motor GPe decreases. For exact definition of ROI within the motor network see table 1. ** = p < 0.005, * = p < 0.05 corrected for multiple comparisons using the network based statistics approach as implemented in the GraphVar toolbox.

In summary, results show that effective STN-DBS leads to an overall increase of connectivity in the motor network of the brain (fig. 3), potentially mediated by a reduced basal ganglia output (and stronger thalamo-cortical interaction, fig. 6). Second, STN-DBS seems to attenuate or “normalize” both functional connectivity profiles seeding from the electrode (fig. 4) and the overall average connectivity (i.e. centrality) estimates in the brain (fig. 5).

## Discussion

Four main conclusions may be drawn from the present study. First, the acquisition of postoperative rs-fMRI in DBS patients is feasible in a clinical setting and resulting data is both useful and useable to noninvasively study effects of DBS on distributed brain networks. Based on functional connectivity estimates to other regions of the motor network, even the BOLD signal sampled directly at and around the electrode exhibited a physiologically meaningful signal. Second, STN-DBS had a significant effect on brain connectivity throughout the motor network, specifically on its cortical and cerebellar subparts. Third, and most importantly, DBS induced changes on average connectivity throughout the motor network were strongly dependent on electrode placement. Specifically, electrodes with strong impact on the motor STN induce larger changes than the ones with weak or no impact on the motor STN (R = 0.7, p < 0.001). This correlation is crucial since it may show a direct modulatory effect of STN-DBS on the motor network. Finally, STN-DBS had the effect of “normalizing” both connectivity profiles of the electrodes but also average connectivity profiles toward profiles found in age-matched healthy control subjects. Again, this effect was dependent on electrode location – well placed electrodes shifted the overall connectivity profiles more strongly toward controls than poorly placed electrodes.

### Clinical implications

The main clinical finding shows that connectivity changes induced by DBS with optimally placed leads are being “normalized” toward healthy controls. Overall, STN-DBS led to increased functional connectivity in the motor network and relative decoupling of basal ganglia pathways. Our findings are based on the largest DBS-fMRI sample to date and extended by explaining individual differences in network changes as a function of DBS electrode placement. This extension is crucial since it shows that fMRI could potentially be used to even guide DBS surgery and programming. Specifically, it could be feasible to acquire real-time rs-fMRI in surgeries that are based on intraoperative MRI. In such a setting, one could picture test stimulations during and after surgery. Lead placement and parameters could then be semi-automatically tuned to maximize impact of DBS on the motor network. Even without intraoperative MRI, before surgery, our findings could be used to guide DBS targeting by estimating the region in and around the STN that is most strongly functionally connected to the motor network. Such a concept has been briefly explored for essential tremor (Anderson *et al.*, 2011) in the past but could be extended to STN-DBS for PD, as well.

### Pathophysiological implications

Beyond clinical implications of our findings, they may also shed light on the pathophysiology of PD and the mechanism of action of DBS. Specifically, we showed an increase of functional connectivity between the motor thalamus and cortex as a function of effective DBS and a decrease of striato-cerebellar coupling. Both effects have partly been shown using fMRI before (Kahan *et al.*, 2014). In particular, the increase of thalamo-cortical interaction by STN-DBS may be derived from classical basal-ganglia cortical loop models (Rodríguez-Oroz *et al.*, 2009). In PD, the dopamine deficit leads to increased activity of the indirect basal ganglia pathway which results in increased GPi/SNr output and in turn inhibition of the ventrolateral thalamic nucleus. STN hyperactivity is a key characteristic of this pathological circuitry and by attenuating this hyperactivity with DBS the inhibition of the thalamo-cortical pathway is reduced (Kahan *et al.*, 2014; Rodríguez-Oroz *et al.*, 2009). Correspondingly, our results suggest an attenuation of coupling between the STN and the striatum as well as the STN and the GPe. Taken the above into account, one may speculate that DBS could “rescue” the thalamocortical interaction due to a disruption of subthalamic control over GABAergic pallidothalamic efferents. In turn, this seems implemented by attenuating coupling between the overactive STN and the striatum. In this way, DBS may restore the hypodopaminergic loss of effective coding capacity within the thalamo-cortical network.

Similarly, our findings suggest that striatal dominance over the cerebellum could be attenuated by effective DBS. Increased functional striato-cerebellar coupling was described in PD patients before (Hou *et al.*, 2016) and attributed to a compensatory effect that was associated to improved movement performance (Simioni *et al.*, 2016). Be it pathologic or compensatory – the increased coupling was decreased to a level found in healthy controls under dopaminergic medication. Our study now extends these findings by showing that STN-DBS seems to have a similar attenuating effect on increased striato-cerebellar coupling.

### DBS as an optimal tool to study the functional connectome of the brain

Our study identified DBS as a promising tool to study changes of the functional connectome due to precise brain stimulation of focal subcortical areas. Currently, such a focal stimulation is not possible using other techniques in the human brain. Moreover, deeper structures are not accessible by noninvasive brain stimulation. World-wide, DBS is an increasingly applied treatment option that is well established for severe movement disorders but indications extend to a growing number of psychiatric diseases (Lozano and Lipsman, 2013). Accordingly, a growing number of structures are targeted by DBS (Fox *et al.*, 2014). Sometimes, the same disease can be treated by targeting varying brain structures. For instance, at least five targets are under investigation to treat obsessive compulsive disorder (de Koning *et al.*, 2011) or treatment-refractory depression (Zhou *et al.*, 2018). Even established diseases are treated with different targets (for instance both PD and dystonia have been treated by stimulating STN, GPi or thalamus). It seems that DBS may modulate symptom-related brain networks that may be overlapping at the same node (e.g. the therapeutic network modulated in PD and dystonia seems to overlap at both STN and GPi). fMRI analyses may help further strengthen this concept. Adding to the complexity, different targets can be used in the same disease to preferentially treat different symptoms (for instance STN being an obvious choice to treat most PD symptoms while the VIM is predominantly used to exclusively treat Parkinsonian tremor). DBS cohorts may now be studied using functional MRI (in DBS ON setting currently approved for some systems by Medtronic only) as was done here and in a few previous studies (table 1). Needless to say, beyond acquiring fMRI data under stimulation in resting-state, changes in task-fMRI paradigms could equally be studied. We argue that DBS-fMRI may soon become a new field of research that may be crucial to investigating the functional architecture of the human brain. Based on this present study, we conclude that much focus should be put upon precise localization of DBS electrode placement. Specifically, studying ON vs. OFF group results without taking electrode placement into account may lead to erroneous conclusions. The reason is that small changes in placement may result in clinically meaningful changes in motor outcome (Horn *et al.*, 2019; Yao *et al.*, 2018) or behavioral response (Irmen *et al.*, 2018; Neumann *et al.*, 2018). In the present study we show that the same is true for functional response patterns.

### Limitations

Several limitations apply for careful interpretation of our results. First and foremost, despite several publications in the past (table 1), the field of fMRI under DBS is still quite new and most publications were case reports or based on low N cohorts of ∼10 patients. Further, the impact of DBS induced artifacts on the rs-fMRI signal has not been investigated in detail and more methodological work is needed to address potential issues, in the future. To our knowledge, several laboratories world-wide are currently investigating methodological and signal-processing questions related to fMRI under DBS. In our study, the BOLD signal sampled from directly around the electrode was at times highly correlated to the SMA (with a maximum R of 0.7 that is unlikely to originate from noise, also see fig. 2), a region that is coupled to the STN and not (directly) impacted by metal artifacts. This made us decide to use the BOLD signal of the electrode itself in one analysis (fig. 4). However, we refrained from using the signal in the most central analyses that draw our main conclusions (figs. 2 & 5). Similarly, the DBS lead extensions typically induce an artifact in cortical structures (as seen in figure 2) that can currently not be avoided. No software algorithms have been introduced to correct for such artifacts given these types of datasets are still comparably new. To reduce the theoretical bias induced by the extension artifact, we chose to use a coarse parcellation of the cortex (while finer parcellations may run into the risk that single parcels are completely filled by the artifact).

A crucial parameter that was investigated here for the first time in an fMRI DBS context was the impact of the stimulation on the motor STN. Several limitations apply for the model used to derive this measure. First, lead reconstruction and patient registration to the STN atlas may be biased. To this end, we use a state-of-the-art DBS imaging pipeline that has been explicitly designed for the task. In fact, a recent study could show that using our pipeline, automatic registrations between patient space and an STN atlas are not significantly different from manual expert segmentations of the nucleus (Ewert, Horn, *et al.*, 2018). The study involved > 11,000 nonlinear warps in > 100 brains and tested 6 multispectral deformation algorithms with several parameter settings each to fit patient data to the STN atlas. Second, the atlas of the STN itself might be biased. To this end, we used a modern atlas that was explicitly created for our pipeline (Ewert, Plettig, *et al.*, 2018) and that is based on the newest multispectral high-resolution MNI template available (Fonov *et al.*, 2009). Recently, others confirmed accuracy of the atlas based on intraoperative microelectrode recordings and showed that it matched electrophysiological data better than three other atlases of the STN (Nowacki *et al.*, 2018). Finally, the motor functional zone of the atlas was defined using diffusion weighted imaging data acquired on specialized MR hardware (Setsompop *et al.*, 2013) and cross-validated using both healthy controls and PD data (Ewert, Plettig, *et al.*, 2018).

A third potential limitation is the impact of the electric field on the STN. In the past, the volume of tissue modulated by DBS (VTA; McIntyre and Grill, 2002) has been modeled as a binary region around the electrode (e.g. Butson *et al.*, 2007; Dembek *et al.*, 2017; Mädler and Coenen, 2012; McIntyre *et al.*, 2004). However, the degree of mesoscopic tissue modulation may not be a binary (“all or nothing”) effect but instead show probabilistic properties – with areas closer to the electrode being more strongly modulated than more distant ones (Eisenstein *et al.*, 2014). In line with this assumption, we were recently able to predict slightly more variance in clinical outcome using a weighted over a binary model (Horn *et al.*, 2019). This led us to adopt the same principle here, i.e. to calculate the sum over the E-field distribution within the motor STN domain. Still, as mentioned earlier, all results shown here would hold significant when repeating the analyses using a binary VTA instead.

Finally, clinical improvement data was not acquired at the time of imaging due to logistic reasons. Surrogate clinical improvement estimates were collected during clinical routine (see results) but the VTAs used during the 12-month visit were substantially different from the VTAs used in the fMRI experiment. Thus, our study describes network effects of DBS on the brain but cannot draw direct conclusions on how those affect clinical outcome. Relatedly, our patients were scanned in the med ON condition for logistical reasons and to reduce additional motion artifacts. Thus, an effect of dopaminergic medication cannot be excluded. However, the two scans directly followed each other (i.e. were performed under comparable medication) and we mainly draw conclusions that are dependent on electrode placement throughout the manuscript. Still, further studies are needed to draw inferences in the clinical and medication domain.

### Conclusions

Our study exemplifies the use of invasive brain stimulation to study and modulate the functional connectome of the human brain. The study shows the promise of using invasive neuromodulation in the exemplary case of PD and hints at the promise to broaden this novel field of functional neuroimaging under precisely targeted and focal brain stimulation. More specifically, we demonstrate that the acquisition of postoperative rs-fMRI under STN-DBS in a clinical setting is feasible and resulting data is useful to noninvasively study DBS effects. DBS may attenuate pathological basal ganglia output leading to changes in average connectivity within the motor network that were strongly dependent on electrode placement. Finally, STN-DBS seemed to have the effect of “normalizing” brain connectivity toward that of healthy controls. Similar studies in populations under DBS with different targets and/or in different diseases will extend our knowledge on the impact of focal brain stimulation on distributed brain networks.

## List of abbreviations

B1+RMS: The root-mean-square value of the MRI Effective Component of the radio frequency (RF) magnetic (B1) field. A measure to control the amount of RF power utilized to assure patient safety (besides B1+RMS, this is often measured by the specific absorption rate; SAR).
BOLD: Blood oxygen level dependent (signal commonly investigated in functional magnetic resonance imaging).
DBS: Deep Brain Stimulation
DCM: Dynamic Causal Modeling
dlPFC: Dorsolateral prefrontal cortex
EC: Eigenvector centrality
GP/GPi/GPe: Globus pallidus / internal part of globus pallidus / external part of globus pallidus
NBM: Nucleus Basalis of Meynert
PD: Parkinson‘s Disease
PDRP: Parkinson‘s Disease related pattern
PET: Positron Emission Tomography
PPN: Pedunculopontine nucleus
UPDRS: Unified Parkinson’s Disease Rating Scale
(rs-)fMRI: (resting-state) functional magnetic resonance imaging
PH: Posterior hypothalamic nucleus
ROI: Region of interest
SMA: Supplementary motor area
STN: Subthalamic nucleus, primary DBS target in PD
VIM: Ventral intermediate nucleus of the thalamus
VP: Ventral posterior nucleus of the thalamus
VPM: Ventral posteromedial nucleus of the thalamus

## Data Availability

The DBS MRI datasets generated during and analyzed during the current study are not publicly available due to data privacy regulations of patient data but are available from the corresponding author on reasonable request. The control cohort is available within the PPMI repository (www.ppmi-info.org). All code used to analyze the dataset is available within Lead-DBS /-Connectome software (https://github.com/leaddbs/leaddbs).

## Acknowledgements

We would like to thank Lea Waller for support on using the GraphVar toolbox. The study was supported by DFG grant SPP 2041 to A.A.K. and DFG Emmy Noether grant 410169619 to A.H. Data used in the preparation of this article were obtained from the Parkinson’s Progression Markers Initiative (PPMI) database (www.ppmi-info.org/data). For up-to-date information on the study, visit www.ppmi-info.org. PPMI – a public-private partnership – is funded by the Michael J. Fox Foundation for Parkinson’s Research and funding partners, see www.ppmi-info.org/fundingpartners.

## Author contributions

A.H. conceptualized the study, developed the software pipeline used, analyzed data and wrote the manuscript. G.W. and J.H. acquired fMRI data and revised the manuscript. F.I. performed the literature analysis and revised the manuscript. N.L. developed the software pipeline used and revised the manuscript. W-J.N., P.K., G.B. and M.S. acquired data, set up DBS compatible MRI sequences and revised the manuscript. A.A.K. conceptualized the study, acquired fMRI data and revised the manuscript.

## Materials & Correspondence

Correspondence and material requests should be addressed to Andreas Horn.

---

**Table S1:**
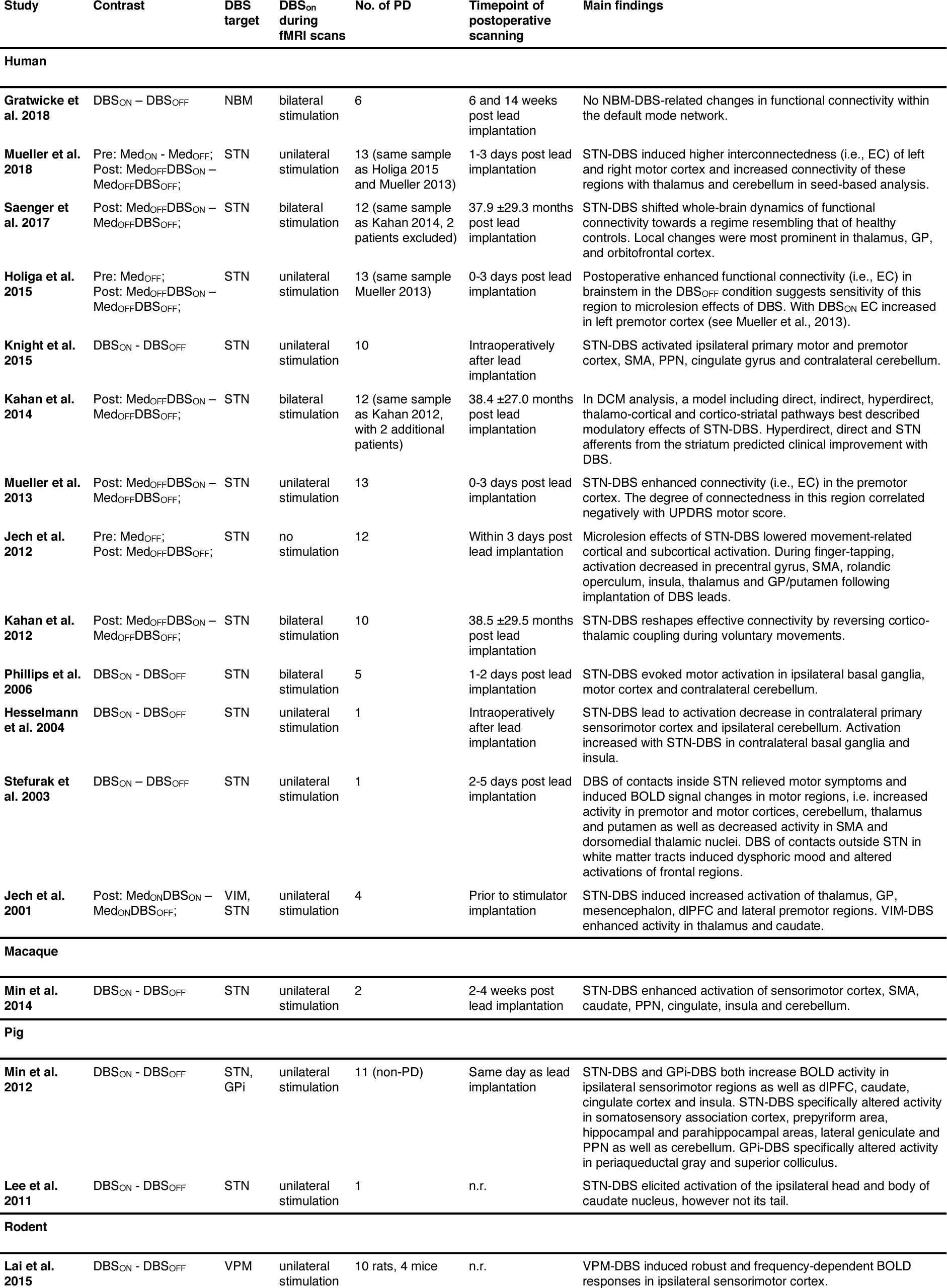
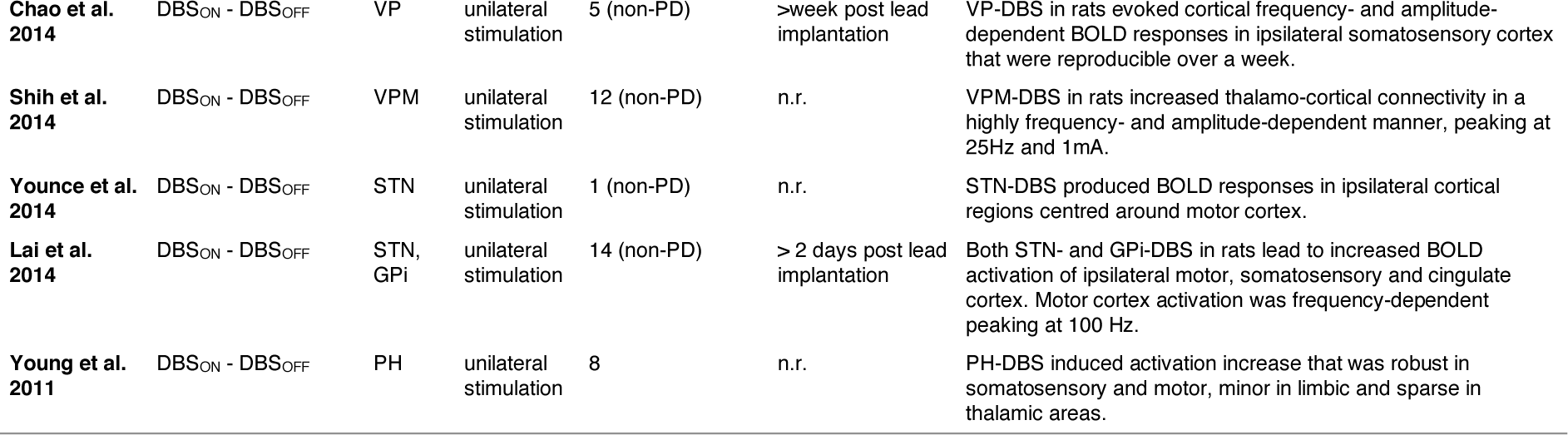
Studies on DBS-induced modulation of resting-state functional connectivity in PD

| Study | Contrast | DBS target | DBS<sub>on</sub> during fMRI scans | No. of PD | Timepoint of postoperative scanning | Main findings |
| --- | --- | --- | --- | --- | --- | --- |
| Human |  |  |  |  |  |  |
| Gratwicke et al. 2018 | DBS<sub>ON</sub> – DBS<sub>OFF</sub> | NBM | bilateral stimulation | 6 | 6 and 14 weeks post lead implantation | No NBM-DBS-related changes in functional connectivity within the default mode network. |
| Mueller et al. 2018 | Pre: Med<sub>ON</sub> - Med<sub>OFF</sub>;<br>Post: Med<sub>OFF</sub>DBS<sub>ON</sub> – Med<sub>OFF</sub>DBS<sub>OFF</sub>; | STN | unilateral stimulation | 13 (same sample as Holiga 2015 and Mueller 2013) | 1-3 days post lead implantation | STN-DBS induced higher interconnectedness (i.e., EC) of left and right motor cortex and increased connectivity of these regions with thalamus and cerebellum in seed-based analysis. |
| Saenger et al. 2017 | Post: Med<sub>OFF</sub>DBS<sub>ON</sub> – Med<sub>OFF</sub>DBS<sub>OFF</sub>; | STN | bilateral stimulation | 12 (same sample as Kahan 2014, 2 patients excluded) | 37.9 ±29.3 months post lead implantation | STN-DBS shifted whole-brain dynamics of functional connectivity towards a regime resembling that of healthy controls. Local changes were most prominent in thalamus, GP, and orbitofrontal cortex. |
| Holiga et al. 2015 | Pre: Med<sub>OFF</sub>;<br>Post: Med<sub>OFF</sub>DBS<sub>ON</sub> – Med<sub>OFF</sub>DBS<sub>OFF</sub>; | STN | unilateral stimulation | 13 (same sample Mueller 2013) | 0-3 days post lead implantation | Postoperative enhanced functional connectivity (i.e., EC) in brainstem in the DBS<sub>OFF</sub> condition suggests sensitivity of this region to microlesion effects of DBS. With DBS<sub>ON</sub> EC increased in left premotor cortex (see Mueller et al., 2013). |
| Knight et al. 2015 | DBS<sub>ON</sub> - DBS<sub>OFF</sub> | STN | unilateral stimulation | 10 | Intraoperatively after lead implantation | STN-DBS activated ipsilateral primary motor and premotor cortex, SMA, PPN, cingulate gyrus and contralateral cerebellum. |
| Kahan et al. 2014 | Post: Med<sub>OFF</sub>DBS<sub>ON</sub> – Med<sub>OFF</sub>DBS<sub>OFF</sub>; | STN | bilateral stimulation | 12 (same sample as Kahan 2012, with 2 additional patients) | 38.4 ±27.0 months post lead implantation | In DCM analysis, a model including direct, indirect, hyperdirect, thalamo-cortical and cortico-striatal pathways best described modulatory effects of STN-DBS. Hyperdirect, direct and STN afferents from the striatum predicted clinical improvement with DBS. |
| Mueller et al. 2013 | Post: Med<sub>OFF</sub>DBS<sub>ON</sub> – Med<sub>OFF</sub>DBS<sub>OFF</sub>; | STN | unilateral stimulation | 13 | 0-3 days post lead implantation | STN-DBS enhanced connectivity (i.e., EC) in the premotor cortex. The degree of connectedness in this region correlated negatively with UPDRS motor score. |
| Jech et al. 2012 | Pre: Med<sub>OFF</sub>;<br>Post: Med<sub>OFF</sub>DBS<sub>OFF</sub>; | STN | no stimulation | 12 | Within 3 days post lead implantation | Microlesion effects of STN-DBS lowered movement-related cortical and subcortical activation. During finger-tapping, activation decreased in precentral gyrus, SMA, rolandic operculum, insula, thalamus and GP/putamen following implantation of DBS leads. |
| Kahan et al. 2012 | Post: Med<sub>OFF</sub>DBS<sub>ON</sub> – Med<sub>OFF</sub>DBS<sub>OFF</sub>; | STN | bilateral stimulation | 10 | 38.5 ±29.5 months post lead implantation | STN-DBS reshapes effective connectivity by reversing cortico-thalamic coupling during voluntary movements. |
| Phillips et al. 2006 | DBS<sub>ON</sub> - DBS<sub>OFF</sub> | STN | bilateral stimulation | 5 | 1-2 days post lead implantation | STN-DBS evoked motor activation in ipsilateral basal ganglia, motor cortex and contralateral cerebellum. |
| Hesselmann et al. 2004 | DBS<sub>ON</sub> - DBS<sub>OFF</sub> | STN | unilateral stimulation | 1 | Intraoperatively after lead implantation | STN-DBS lead to activation decrease in contralateral primary sensorimotor cortex and ipsilateral cerebellum. Activation increased with STN-DBS in contralateral basal ganglia and insula. |
| Stefurak et al. 2003 | DBS<sub>ON</sub> – DBS<sub>OFF</sub> | STN | unilateral stimulation | 1 | 2-5 days post lead implantation | DBS of contacts inside STN relieved motor symptoms and induced BOLD signal changes in motor regions, i.e. increased activity in premotor and motor cortices, cerebellum, thalamus and putamen as well as decreased activity in SMA and dorsomedial thalamic nuclei. DBS of contacts outside STN in white matter tracts induced dysphoric mood and altered activations of frontal regions. |
| Jech et al. 2001 | Post: Med<sub>ON</sub>DBS<sub>ON</sub> – Med<sub>ON</sub>DBS<sub>OFF</sub>; | VIM, STN | unilateral stimulation | 4 | Prior to stimulator implantation | STN-DBS induced increased activation of thalamus, GP, mesencephalon, dIPFC and lateral premotor regions. VIM-DBS enhanced activity in thalamus and caudate. |
| Macaque |  |  |  |  |  |  |
| Min et al. 2014 | DBS<sub>ON</sub> - DBS<sub>OFF</sub> | STN | unilateral stimulation | 2 | 2-4 weeks post lead implantation | STN-DBS enhanced activation of sensorimotor cortex, SMA, caudate, PPN, cingulate, insula and cerebellum. |
| Pig |  |  |  |  |  |  |
| Min et al. 2012 | DBS<sub>ON</sub> - DBS<sub>OFF</sub> | STN, GPI | unilateral stimulation | 11 (non-PD) | Same day as lead implantation | STN-DBS and GPI-DBS both increase BOLD activity in ipsilateral sensorimotor regions as well as dIPFC, caudate, cingulate cortex and insula. STN-DBS specifically altered activity in somatosensory association cortex, prepyriform area, hippocampal and parahippocampal areas, lateral geniculate and PPN as well as cerebellum. GPI-DBS specifically altered activity in periaqueductal gray and superior colliculus. |
| Lee et al. 2011 | DBS<sub>ON</sub> - DBS<sub>OFF</sub> | STN | unilateral stimulation | 1 | n.r. | STN-DBS elicited activation of the ipsilateral head and body of caudate nucleus, however not its tail. |
| Rodent |  |  |  |  |  |  |
| Lai et al. 2015 | DBS<sub>ON</sub> - DBS<sub>OFF</sub> | VPM | unilateral stimulation | 10 rats, 4 mice | n.r. | VPM-DBS induced robust and frequency-dependent BOLD responses in ipsilateral sensorimotor cortex. |
| **Chao et al. 2014** | DBS<sub>ON</sub> - DBS<sub>OFF</sub> | VP | unilateral stimulation | 5 (non-PD) | >week post lead implantation | VP-DBS in rats evoked cortical frequency- and amplitude-dependent BOLD responses in ipsilateral somatosensory cortex that were reproducible over a week. |
| **Shih et al. 2014** | DBS<sub>ON</sub> - DBS<sub>OFF</sub> | VPM | unilateral stimulation | 12 (non-PD) | n.r. | VPM-DBS in rats increased thalamo-cortical connectivity in a highly frequency- and amplitude-dependent manner, peaking at 25Hz and 1mA. |
| **Younce et al. 2014** | DBS<sub>ON</sub> - DBS<sub>OFF</sub> | STN | unilateral stimulation | 1 (non-PD) | n.r. | STN-DBS produced BOLD responses in ipsilateral cortical regions centred around motor cortex. |
| **Lai et al. 2014** | DBS<sub>ON</sub> - DBS<sub>OFF</sub> | STN, GPi | unilateral stimulation | 14 (non-PD) | > 2 days post lead implantation | Both STN- and GPi-DBS in rats lead to increased BOLD activation of ipsilateral motor, somatosensory and cingulate cortex. Motor cortex activation was frequency-dependent peaking at 100 Hz. |
| **Young et al. 2011** | DBS<sub>ON</sub> - DBS<sub>OFF</sub> | PH | unilateral stimulation | 8 | n.r. | PH-DBS induced activation increase that was robust in somatosensory and motor, minor in limbic and sparse in thalamic areas. |

